# Pumilio proteins license viral noncoding RNA engagement with target mRNAs

**DOI:** 10.64898/2026.10.06.757068

**Authors:** Carlos Gorbea, Demián Cazalla

**Affiliations:** Department of Biochemistry, University of Utah, Salt Lake City, UT, USA

## Abstract

Noncoding RNAs commonly rely on RNA-binding proteins (RBPs) to execute their regulatory functions, but whether RBPs can instead facilitate target recognition by noncoding RNAs is poorly understood. Here, we identify a non-canonical function for Pumilio proteins PUM1 and PUM2 in gene regulation as cofactors of the viral small nuclear RNAs HSUR1 and HSUR2 encoded by *Herpesvirus saimiri*. HSUR1 and HSUR2 function as microRNA and RBP adaptors that bind target mRNAs to downregulate their expression. We show that HSUR1 and HSUR2 contain conserved Pumilio response elements (PREs) through which they associate with PUM1 and PUM2, without measurably altering PUM protein activity or subcellular distribution in infected cells. Instead, PUM binding is required for HSUR-mediated mRNA repression. Mutation of the PREs abolishes PUM association and HSUR-mediated repression without disrupting HSUR association with microRNAs, whereas tethering PUM1 or PUM2 directly to PRE-deficient HSURs restores repression. Furthermore, HSURs unable to bind PUMs fail to associate with target mRNAs, demonstrating that PUM binding is required for productive HSUR-mRNA engagement. These findings reveal that PUM proteins can function as enabling factors for noncoding RNA target recognition rather than as downstream effectors of repression, expanding the functions of these RBPs in RNA-mediated gene regulation.

## INTRODUCTION

In recent decades, noncoding RNAs (ncRNAs) have emerged as pervasive regulators of gene expression. NcRNA-mediated regulation generally requires interactions with cellular RNAs and proteins. In most cases, RNA binding proteins (RBPs) that associate with ncRNAs function as downstream effectors, executing the regulatory outcomes specified by the ncRNAs. For example, in canonical microRNA (miRNA)-mediated gene silencing, miRNAs function primarily as sequence-specific guides that direct Argonaute (Ago) proteins to complementary sites within target RNAs. Once recruited to the target, Ago serves as the principal effector of gene silencing by engaging TNCR6 adaptor proteins, which in turn recruit deadenylase complexes, including the CCR4-NOT complex, and other inhibitory factors to promote transcript destabilization and/or translational repression (1). The coupling of RNA-guided target recognition with protein-mediated execution is a broadly established paradigm for ncRNA function.

In some cases, ncRNAs can also recruit other ncRNAs to target RNAs. HSUR1 and HSUR2 are two small nuclear RNAs (snRNAs) of the Sm-class (2) expressed by the oncogenic *Herpesvirus saimiri*(3) (HVS) in latently infected cells. Both HSUR1 and HSUR2 function as miRNA adaptors that bind host miRNAs miR-142-3p and miR-16 (4) and recruit them to 3′ untranslated regions (3′UTRs) of target mRNAs to repress their expression (5–7). HSUR1 regulates genes involved in RNA metabolism, chromatin organization and transcription, thereby acting as master coordinator of gene expression that influences a large fraction of the transcriptome of HVS-infected cells (7). HSUR1 also binds miR-27 to trigger its degradation through target-directed miRNA decay (4,8). In addition to miR-142-3p, HSUR1 can also recruit AU-rich (ARE)-binding proteins (ARE-BPs) to target mRNAs, further enhancing HSUR1-mediated mRNA repression (7). Binding of miR-142-3p to the 5′ end of HSUR1 and HSUR2 (Fig. 1A) is required for target RNA repression and recruitment of ARE-BPs and miR-16, respectively (5,7,9). Accordingly, disruption of the miR-142-3p interaction renders both HSUR1 and HSUR2 unable to repress target mRNAs (7,9).

**Figure 1.**
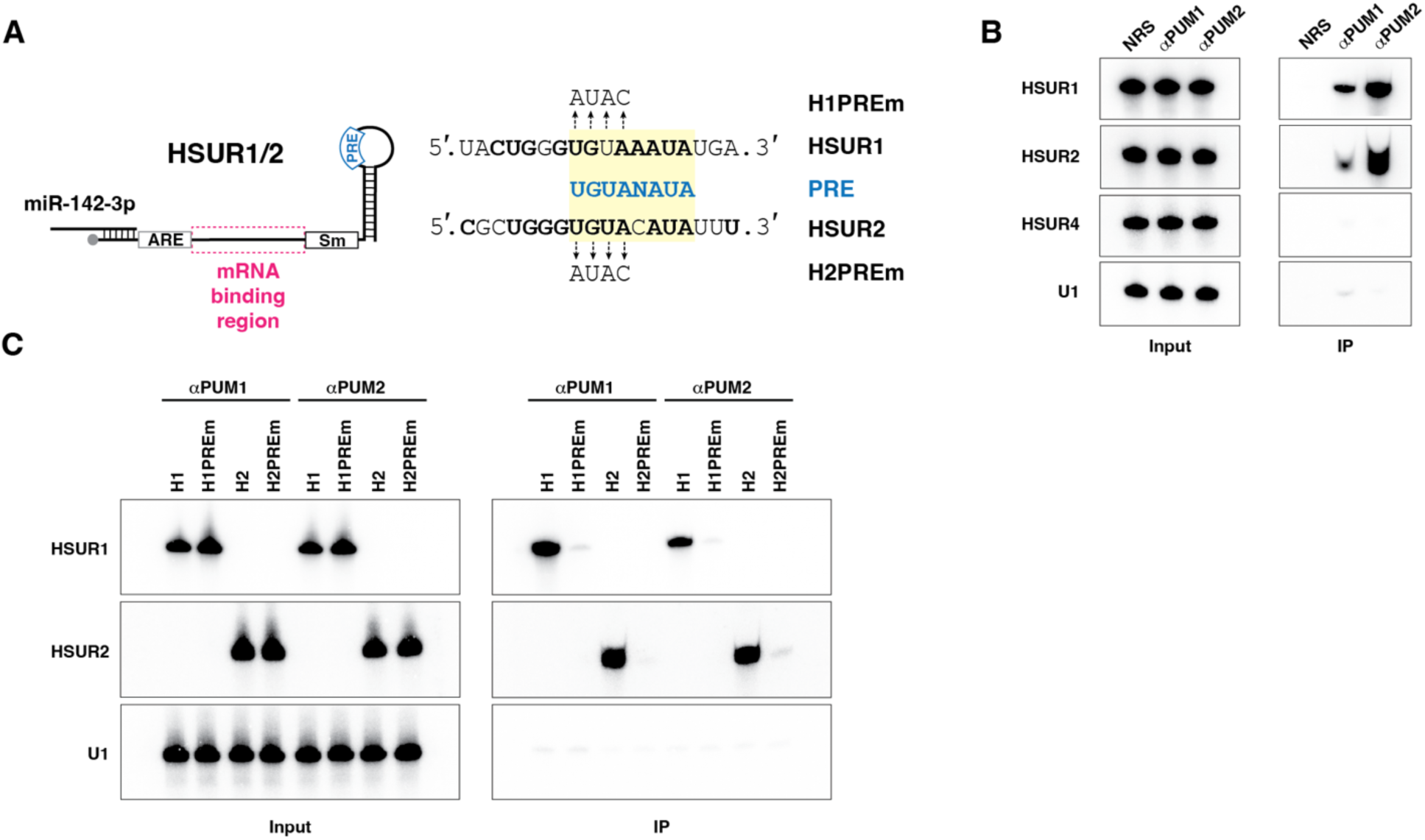
HSUR1 and HSUR2 bind PUM proteins in HVS-infected cells. **(A)** Schematic representation of HSUR1 or HSUR2 (HSUR1/2). The locations of the miR-142-3p binding site, the region involved in mRNA binding, and Pumilio response element (PRE) are indicated. The sequences within the 3′ loops of HSUR1 and HSUR2 that constitute the PREs, as well as the mutations introduced into these PREs used to generate H1PREm and H2PREm, are shown on the right. Sm, Sm-binding site; PRE, Pumilio response element; ARE, AU-rich element. **(B)** Co-immunoprecipitation of HSUR1 and HSUR2 with normal rabbit serum (NRS) or antibodies against PUM1 and PUM2 from cj319-WT cells. Northern blotting was used to detect HSUR1, HSUR2, as well as HSUR4 and cellular U1 snRNA as immunoprecipitation controls. Input, 5%; IP, 95%. **(C)** As in (B) from extracts prepared from HeLa cells expressing either HSUR1, HSUR2, H1PREm or H2PREm.

Unlike miRNAs, HSUR1and HSUR2 do not contain a unique seed sequence that mediate interactions with target mRNAs. Instead, both ncRNAs base-pair with mRNAs through variable stretches of nucleotides within an extended central domain of the snRNA (Fig. 1A), with distinct binding sites exhibiting unique base-pairing configurations. The mechanism by which HSUR1 and HSUR2 can specifically recognize their binding sites in target mRNAs remains unknown. The absence of a defined seed region suggests that sequence complementarity between HSUR1 and HSUR2 and target mRNAs is required, but may not be sufficient, for the selection of binding sites. It has been proposed that, analogous to cellular spliceosomal snRNAs that rely on *cis-*acting signals like splicing enhancers and *trans*-acting factors such as SR proteins to guide and escort spliceosomal small nuclear ribonucleoprotein particles (snRNPs) to splice sites (10–15), HSURs may similarly require RBPs to assist in the recognition of binding sites in target mRNAs (6).

The Pumilio (PUM) family of RNA binding proteins binds target mRNAs and promotes their deadenylation by directly recruiting the CCR4-NOT deadenylase complex (16,17). The two human paralogs, PUM1 and PUM2, exhibit the same repressive activities by binding with high affinity to a specific Pumilio Response Element (PRE) with the consensus 5′-UGUANAUA-3′ (N=U,A,G,C) sequence (18). PUM proteins have conserved functions in development and neurological processes, and their dysfunction can contribute to neurological disorders and cancer (19). PUM proteins are predominantly cytoplasmic and can be enriched in processing bodies (P-bodies) and stress granules (20–22). The activities of PUM1 and PUM2 can also be modulated by the abundant, cytoplasmic long ncRNA called *NORAD* (ncRNA activated by DNA damage) (23,24). *NORAD* contains ∼17-20 PREs and acts as a competitive inhibitor by sequestering PUM proteins, thereby increasing the expression of PRE-containing mRNAs (23).

Interactions between HSUR1 and HSUR2 and host miRNAs were originally suggested by analyses of sequence conservation among HVS strains and the related *Herpesvirus ateles* (HVA) (4). These analyses revealed perfectly or highly conserved sequences corresponding to binding sites for miR-142-3p, miR-16 and miR-27, as well as AREs and binding sites for Sm-proteins. In addition, they identified a highly conserved motif of unknown function located within the 3′ loop of both HSUR1 and HSUR2 (Fig. 1A). In this study, we show that this conserved motif constitutes a PRE and that HSUR1 and HSUR2 associate with PUM1 and PUM2 in HVS-infected cells. We further show that, unlike *NORAD*, HSUR1 and HSUR2 do not affect PUM1 and PUM2 activities in HVS-infected cells. Instead, binding of PUM proteins to the PRE present within these HSURs is required for HSUR1- and HSUR2-mediated mRNA repression.

Furthermore, our data suggest that HSUR-bound PUM proteins do not function as downstream effectors that promote deadenylation of target mRNAs. Rather, PUM binding to HSUR1 and HSUR2 enables their association with target mRNAs, revealing a non-canonical role for PUM proteins in ncRNA-mediated mRNA repression.

## MATERIALS AND METHODS

### Cell and virus culture, transfections

Cj319-WT and cj319-Δ2a cell lines were grown as previously described (5). 293T/17 (CRL 11268) and HeLa cells (CCL-2) were purchased from ATCC and grown as described (9). HuT78 Sézary Syndrome, cutaneous T lymphocytes (obtained from Vicente Planelles) were grown as described (7). Cell lines were not authenticated. All cell lines were routinely tested for the presence of mycoplasma using the universal mycoplasma detection kit from ATCC.

For transient knockdown of HSUR1 and HSUR2, 10 million cj319-WT marmoset T cells were nucleofected in a nucleofector™ 2b device as described previously (5) with 1 nmol of chimeric antisense oligonucleotide (ASO; Integrated DNA Technologies) targeting HSUR1 (7) and 0.5 nmol of ASO targeting HSUR2 (5), or 1.5 nmol of ASO targeting the mRNA encoding the green fluorescent protein (GFP) (5) as a control. All ASOs contained a phosphorothioate backbone and five 2’-*O*-methoxyethyl ribonucleotides at each end. The sequences of the ASOs are included in Supplementary Table 1. These ASOs specifically induced degradation of HSUR1 and HSUR2. Twenty-one hours after transfection the cells were harvested, washed with PBS and stored in TRIzol® at -80°C until used. For transient knockdown of PUM1 and PUM2, ten million cj319-WT or cj319-Δ2a marmoset T cells were similarly nucleofected in a nucleofector™ 2b device with 100 pmoles each of siRNAs targeting PUM1 (Silencer Select siRNA assay s18680, Thermo Fisher Scientific) and PUM2 (Silencer Select siRNA assay s23671, Thermo Fisher Scientific) or 200 pmoles of siRNAs targeting GFP (Integrated DNA Technologies) as control. All siRNA sequences can be found in Supplementary Table 1.

All lentiviral transductions were performed in HuT78 or HeLa cells using supernatants generated as previously described(9). Stable cell lines were generated by selection with blasticidin for *Renilla* luciferase-expressing HuT78 cells; with blasticidin and puromycin for *Renilla* and firefly luciferase-expressing HuT78 cells; or with puromycin alone for HeLa cells expressing HSUR1, HSUR2, HSUR1 carrying mutations in the PRE sequence (H1PREm) or HSUR2 carrying mutations in the PRE sequence (H2PREm). U937 cells expressing luciferase reporters sensitive to HSUR2 have been described (6). Stably transduced HuT78 and U937 cells expressing firefly and *Renilla*-luciferase were grown in complete medium plus 8 µg/ml of puromycin (InvivoGen) and 10 µg/ml of blasticidin (InvivoGen). Stably transduced HeLa cells expressing wild-type or mutant versions of HSUR1 or HSUR2 were grown in complete medium supplemented with 10 µg/ml or 8 µg/ml of puromycin, respectively.

HeLa cells were transfected as previously described (12) with Lipofectamine RNAiMax (Thermo Fisher Scientific) according to the manufacturer’s instructions. Cells in 10-cm plates were transfected with 150 pmol of either Control miRNA or miR-142-3p (see Supplementary Table 1 for sequences). Cells grown in six-well plates were transfected with 25 pmol of miRNA as described previously (9). In all experiments, cells were harvested approximately 24 h after transfection.

### Plasmids

Plasmids for stable expression of HSUR1, HSUR2, and firefly and *Renilla* luciferase reporters have been already described (6,7,9). Plasmids for transient expression of HSUR1 or HSUR2 and their respective control plasmids have also been described (6,7). Plasmids encoding PRE-mutant versions of HSUR1 and HSUR2 for either constitutive or transient expression were generated by site-directed mutagenesis. Plasmid for transient expression of HSUR4 has been described (25). Plasmids for transient expression of mutant versions of HSUR1 and HSUR2 in which the 3′ stem-loops were replaced with boxB hairpins were generated by Gibson assembly (26) of the backbone of plasmids encoding HSUR1 or HSUR2 and DNA fragments containing the boxB hairpin sequence (Integrated DNA Technologies) with appropriate Gibson overlaps. Plasmids for transient expression of λN-PUM1 or λN-PUM2 were generated by Gibson assembly using the backbone of ZFP36L2-mCherry2 (7) and DNA fragments encoding λN-PUM1 or λN-PUM2 (Twist Biosciences) with appropriate Gibson overlaps. All plasmids generated by Gibson assembly were fully sequenced Plasmidosaurus (San Francisco, CA).

To generate plasmids expressing firefly luciferase reporters with MS2 hairpins, fragments containing the MS2 hairpins were amplified by PCR using the plasmid pSL-MS2-12X (Addgene #27119) (27) as template, and inserted into a unique *Sal*I site in the plasmid pLenti-TK-Firefly-STK4 (6) or pLenti-TK-Firefly-JARID2_BS1 (7). The insertion and orientation of the MS2 hairpin fragments were confirmed by sequencing. The plasmid pMS2-GFP (28) for transient expression of enhanced GFP (eGFP) fused to the MS2 coat protein (MCP), was obtained from Addgene (#27121).

### Preparation of nuclear and cytoplasmic fractions from marmoset cj319-WT and cj319-Δ2a cells

Cells were counted, harvested, centrifuged and washed once with cold phosphate-buffered saline (PBS), pH 7.5. Duplicate samples containing 1x10^7^ cj319-WT or cj319-Δ2a cells were each suspended in 100 µl of extraction buffer (50 mM Tris-HCl, pH 7.5, 0.32 M sucrose, 3 mM CaCl_2_, 2 mM magnesium acetate, 1 mM DTT, SUPERase•In^®^ RNase inhibitor (5 µl per 100 µl of extraction buffer; Thermo Fisher Scientific), cOmplete™ protease inhibitors (one tablet per 10 ml of extraction buffer; Roche), 1 mM PMSF (Sigma-Aldrich) and 0.25% of Nonidet P-40 (NP-40) substitute (Roche) by pipetting followed by vortexing. Cells were incubated on ice for 5 min with occasional vortexing every 1-2 min. Lysates were centrifuged at 500 x *g* for 5 min at 4°C, and the supernatant fractions (cytoplasmic fractions) were transferred to fresh tubes and kept on ice. One replicate supernatant fraction from each cell line was immediately mixed with 1 ml of TRIzol^®^ (Thermo Fisher Scientific) and stored at -80°C until use, whereas the second replicate supernatant fraction from each cell line was mixed with 100 µl of 4X sodium dedoceyl sulphate-polyacrylamide gel electrophoresis (SDS-PAGE) sample buffer, vortexed and incubated on a heating block at 95°C for 5 min. The pellets from each extraction were then washed with extraction buffer lacking NP-40 and centrifuged. The replicate pellets (nuclear fractions) from each cell line were suspended in 100 µl of wash buffer and immediately mixed with 1 ml of TRIzol^®^ or 100 µl of 4X SDS-PAGE sample buffer as described above. Total fractions from cj319-WT cells and cj319-Δ2a cells were prepared from duplicate samples containing 1x10^7^ cells each, which were suspended in 100 µl of wash buffer and immediately mixed with either 1 ml of TRIzol^®^ or 100 µl of 4X SDS-PAGE sample buffer.

### Northern and immunoblotting of nuclear and cytoplasmic fractions from cj319-WT and cj319-Δ2a cells

For Northern blotting, total RNA was separated on 8% denaturing polyacrylamide gels for snRNA and transfer RNA (tRNA) detection and on 14% gels for miRNA and snRNA detection. Northern blots were performed as previously described (5) using DNA oligoprobes described in Supplementary Table 1. Samples in 2X SDS-PAGE sample buffer were incubated at 95°C for 5 min, with nuclear fractions briefly sonicated to shear DNA and reduce viscosity. Twenty-microliter aliquots of each fraction were separated on 4-20% gradient gels (Bio-Rad) using high Tris running buffer (50 mM Tris base, 193 mM glycine, 0.1% SDS). Proteins were transferred onto nitrocellulose membranes and probed with rabbit polyclonal antibodies against PUM1 (1:2000 dilution; Proteintech, Cat. No. 26256-1-AP), a rabbit polyclonal antibody against PUM2 (1:500 dilution; Bethyl-Fortis Life Sciences, Cat. No. A300-202A), or with rabbit monoclonal antibodies against GAPDH (1:5000 dilution; Cell Signaling Technology, clone 14C10, Cat. No. 2118) or Lamin B1 (1:1000 dilution; Cell Signaling Technology, clone D4Q4Z, Cat. No. 12586).

Proteins were visualized with IR-Dye^®^ 680RD goat anti-rabbit IgG secondary antibodies and an Odyssey fluorescent scanner (LI-COR). The relative abundance of each protein in the cytoplasmic and nuclear fractions was determined by densitometry using ImageJ2 version 2.16.0/1.54p and expressed as a fraction of the amount of each protein in the total extracts cj319-WT or cj319-Δ2a cells. Data were graphed with Prism 10 software, and statistical significance was determined by two-way ANOVA analysis followed by Tukey’s multiple comparison test (Supplementary Table 2).

### Expression of HSUR1 or HSUR2 bearing mutations at the PRE sequence

Two million HuT78 cells expressing reporter constructs consisting of firefly luciferase fused to 3’UTR sequences from HNRNPA2B1_BS2 or JARID2_BS1 (HSUR1 targets) (7) or 2x10^6^ U937 cells expressing reporter constructs from TP53RK or STK4 (HSUR2 targets) (6) were electroporated with 7.5 µg of control plasmids encoding GFP together with either the U1 promoter (for comparison with HSUR1-expressing plasmids) (7) or the HSUR2 promoter (for comparison with HSUR2-expressing plasmids) (6), or 7.5 µg of plasmids encoding GFP together with either wild-type HSUR1 (7) or H1PREm under control of the U1 promoter, or wild-type HSUR2 (6) or H2PREm under the control of its own promoter. Electroporation was performed using a Neon device by resuspending cells in 100 µl of buffer T containing the indicated plasmids and applying three 1,500-V pulses of 10 ms each for HuT78 cells or three 1,900-V pulses of 10 ms each for U937 cells. Electroporated cells were then cultured for approximately 21 hours in 2 ml of complete RPMI-1640 medium containing 10 µg/ml each of blasticidin and puromycin. Cells were harvested, centrifuged and treated with Stem-Pro Accutase^®^ (Thermo Fisher Scientific) for 3 min. Cells were recovered by centrifugation and resuspended in 0.5 ml of complete media containing 100 ng/ml of 2-(4-amidinophenyl)-1H-indole-6-carboxamidine (DAPI). Thirty thousand DAPI-negative, GFP-positive cells were sorted by fluorescence-activated cell sorting (FACS) using a BD Aria cell sorter equipped with 408-nm violet (450/50 bandpass filter) and 488-nm blue (530/30 bandpass filter) lasers, FACSDiva 8.0 software and an 85-µm nozzle. Sorted cells were collected in Eppendorf tubes containing 0.5 ml of culture media.

### Luciferase reporter assays

Sorted cells were incubated at 37°C with 5% CO_2_ for 30 minutes and centrifuged at 1,000 X *g* for 20 min at room temperature. The supernatant was aspirated, and cells were lysed with 50 µl of 1X Passive Lysis Buffer (Promega) for 20 min at room temperature. Firefly luciferase activity was measured in triplicate using 10 µl of cell lysate per well in a 96-well plate and 50 µl of Luciferase Assay Reagent II from the Dual-Luciferase^®^ Reporter Assay System (Promega) on a Modulus II microplate reader with an integration time of 5 s. Immediately thereafter, 50 µl of STOP & GLO^®^ reagent were added to each well, and *Renilla* luciferase activity was measured using the same parameters. Data were graphed with Prism 10 software. Statistical significance was determined separately for each reporter using ordinary one-way ANOVA, which indicated significant difference among the means for HNRNPA2B1 (*P*<0.0001), JARID2_BS1 (*P*=0.0004), STK4 (*P*<0.0001), and TP53RK (*P*=0.0029), followed by an uncorrected Fisher’s least significant difference (LSD) multiple-comparison test (Supplementary Table 2).

### Quantitative real-time polymerase chain reaction (qRT-PCR)

Total RNA was purified from cells stored in TRIzol^®^ according to the manufacturer’s protocol. Samples included 1x10^7^ cj319-WT marmoset T cells transfected with GFP ASO or a mixture of HSUR1 and HSUR2 ASOs; 1x10^7^ cj319-WT or cj319-Δ2a cells transfected with either control siRNA or a mixture of two siRNAs targeting PUM1 and PUM2; and HeLa cells expressing wild-type HSUR1 or HSUR2 or HSURs bearing mutations in the PRE sequence that were transfected with either control miRNA or miR-142-3p. Plasmids for stable expression of HSUR1 and firefly and *Renilla* luciferase reporters have been previously described (6,9). Following the isopropanol precipitation step, RNA was suspended in 85 µl of water, mixed with 10 µl of 10X DNase I buffer (New England Biolabs) and 5 µl of RNase-free DNase I (New England Biolabs), and incubated at 37°C for 30 min. RNA was purified on RNA Clean & Concentrator™-5 columns with two volumes of RNA binding buffer and one volume of 100% ethanol according to the manufacturer’s instructions (Zymo Research). cDNA was synthesized in 20-µl reactions from 900 ng of DNase I-treated total RNA using the High-Capacity cDNA Reverse Transcription Kit with MultiScribe Reverse Transcriptase and random primers (Applied Biosystems). Real-time PCR reactions were performed in triplicate for each independent experiment in 8-µl reactions using primers at 0.5 µM, cDNA diluted 1:10 (except for ribosomal 18S, for which cDNA was diluted 1:50,000), and PerfectCTa^®^SYBR^®^Green FastMix^®^ (Quantabio) in a Bio-Rad CFX Opus 384 Real Time PCR system. The cycling conditions were 95°C for 5 min for one cycle, followed by 55 cycles of 95°C for 10 s, 60°C for 10 s and 72°C for 10 s, followed by a melting-curve analysis from 65°C to 95°C with a ramp rate of 0.5°C/s and a 5-s plate read). qPCR primers (Supplementary Table 1) were designed using Primer3Plus (https://www.primer3plus.com/index.html) and validated using cDNA prepared from 900 ng of DNAse I-treated total RNA from each cell line in 20-µl reactions and serially diluted 1:5 to 1:50,000. *C*t values were determined using the instrument’s automated baseline and threshold settings. Relative abundance of HSUR1, HSUR2 or PUM target mRNAs was calculated using the 2^-ΔΔ*C*t^ method by dividing the 2^-Δ*C*t^ value for each target in the treated sample (HSUR1/HSUR2 ASO, PUM1/PUM2 siRNA, or miR-142-3p) by the corresponding 2^-Δ*C*t^ in the control sample (control ASO, control siRNA or control miRNA). For normalization, the *C*t value for 18S ribosomal RNA was subtracted from the *C*t value of each HSUR1, HSUR2 or PUM targets to calculate Δ*C*t. Data were graphed with Prism 10 software and statistical significance was determined by multiple uncorrected Student’s *t*-tests (Supplementary Table 2)

### Co-expression of boxB-containing HSUR1 or HSUR2 with λN-PUM1 or λN-PUM2

Two million HuT78 cells expressing firefly luciferase fused to 3’UTR sequences from HNRNPA2B1 or JARID2 were electroporated with a plasmid mixture containing 6.5 µg of control plasmid encoding GFP and the U1 promoter, or 6.5 µg of a plasmid encoding U1 promoter and a mutant version of HSUR1 in which 3′ terminal stem-loop was replaced with a boxB hairpin.

Each mixture also contained 1 µg of either Control-mCherry2 plasmid encoding mCherry alone, or 1 µg of plasmid encoding mCherry and either λN-PUM1 or λN-PUM2. Electroporation was carried out in a Neon device as described above.

For HSUR2 reporters, 2x10^6^ U937 cells expressing firefly luciferase fused to 3’UTR sequences of STK4 or TP53RK were electroporated with a plasmid mixture containing either 6.5 µg of a control plasmid encoding GFP and HSUR2 promoter plus 1 µg of plasmid encoding mCherry, or 6.5 µg of plasmid encoding wild-type HSUR2 or HSUR2*boxB* mutant plus 1 µg of plasmid encoding mCherry with λN-PUM1 or λN-PUM2. Cells were electroporated using the Neon device as described above. Following electroporation, HuT78 and U937 cells were grown for 21 hours in 2 ml of complete RPMI-1640 medium containing 10 µg/ml each of blasticidin and puromycin. Cells were harvested and suspended in 0.5 ml of complete medium containing DAPI as described above. Thirty thousand DAPI-negative, GFP- and mCherry-double positive cells were sorted by FACS using a BD Aria cell sorter equipped with 408-nm violet (450/50 bandpass filter), 488-nm blue (530/30 bandpass filter) and 561-nm yellow-green (615/25 bandpass filter) lasers and collected in Eppendorf tubes containing 0.5 ml of culture medium. Firefly and *Renilla* luciferase activities were measured in the sorted cell lysates as described above. Data were graphed with Prism 10 software. Statistical significance was determined separately for each reporter using ordinary one-way ANOVA, which indicated significant difference among the means for HNRNPA2B1 (*P*=0.0019), JARID2_BS1 (*P*<0.0001), STK4 (*P*=0.0006), and TP53RK (*P*,0.0029), followed by an uncorrected Fisher’s least significant difference (LSD) multiple-comparison test (Supplementary Table 2).

### Measurement of HSUR1 and HSUR2 association with target reporter transcripts containing MS2 hairpins

For each transfection experiment, a master mix was prepared by mixing 300 µl of buffer T for the Neon device with 3 µg of plasmid encoding GFP fused to MCP and 19.5 µg of either control plasmid, a plasmid encoding wild-type HSUR1 or HSUR2, or plasmids encoding mutant versions of HSUR1 or HSUR2 that do not bind miR-142-3p (H1m142 and H2m142) (9), PUM proteins (H1PREm and H2PREm), or the 3’UTR sequence fused to firefly luciferase expressed by the reporter lines (JARID2_BS1 in the case of HSUR1, STK4 in the case of HSUR2). For HSUR1, 2x10^6^ HuT78 cells expressing firefly luciferase fused to JARID2 3’UTR sequence linked to 10 MS2 hairpin repeats were electroporated in triplicate with 100 µl of master mix containing HSUR1, H1m142, H1PREm or H1mJARID2BS1, as described above. For HSUR2, 2x10^6^ HuT78 cells expressing firefly luciferase fused to STK4 3’UTR sequence linked to 6 MS2 hairpin repeats were electroporated in triplicate with a master mix containing HSUR2, H2m142, H2PREm, or H2mSTK4, as described above. The electroporated cells were pooled and cultured in 5 ml of complete culture medium for 21 hours at 37°C with 5% CO_2_. Cells were then harvested, washed with PBS, and collected by centrifugation.

Cell pellets were suspended on ice in 250 µl of NET2 buffer (100 mM Tris-HCl, pH 7.5, 150 mM NaCl, 0.5% NP-40 substitute [Roche]) containing 100 U of SUPERase•In^®^ RNase inhibitor, cOmplete™ protease inhibitors (one tablet for each 5 ml) and 1 mM PMSF. Cell extracts were prepared by sonication using a Branson 450 sonicator with ten pulses at output 3 and 50% duty cycle. Ten percent of each extract was saved in 1 ml of TRIzol^®^ as input fractions and stored at -80°C until used. The remaining extracts were incubated for 5 hours at 4°C with continuous rotation with 25 µl of protein A Dynabeads™ (Thermo Fisher Scientific) coupled to 2.5 µg of rabbit polyclonal antibodies to GFP (Proteintech). Beads were collected using a magnetic rack and washed three times with cold NET2 buffer. After the third wash, beads were transferred to a fresh Eppendorf tube and stored with 1 ml of TRIzol^®^ at -80°C until used.

Total RNA was purified from the input and anti-GFP immunoprecipitated fractions in TRIzol^®^ according to the manufacturer’s protocol. Three microliters of each input fraction and 14.2 µl of each immunoprecipitated fraction were used to prepare cDNA in 20-µl reactions as described above. Real-time PCR was performed in 8-µl reactions using cDNA diluted 1:2, primers at a concentration of 0.5 µM and PerfectCTa^®^SYBR^®^Green FastMix^®^ in a Bio-Rad CFX opus 384 real-time PCR system as described above. qPCR primers for the firefly luciferase gene expressed in the reporter HuT78 cell lines were designed using Primer3Plus (https://www.primer3plus.com/index.html) and validated using cDNA prepared from 900 ng of DNase I-treated total RNA from each reporter line in 20-µl reactions and serially diluted 1:5 to1:50,000. Primers for HSUR1 and HSUR2 were designed empirically by choosing sequences immediately upstream of the PRE sequence and upstream of the sequences in HSUR1 or HSUR2 engaged in base-pairing with target sequences in the luciferase-JARID2_BS1 or the luciferase-STK4 reporters, respectively. An online Tm calculator from Thermo Fisher Scientific was used to ensure an annealing temperature of at least 60°C for each primer pair. Primers were validated using cDNA prepared from 900 ng of DNase I-treated total RNA from HeLa cells lines expressing HSUR1 or HSUR2 and serially diluted 1:5 to 1:50,000. *C*t values were determined using the instrument’s automated functions for baseline and threshold setting and *C*t determination.

The relative abundance of HSUR1 or HSUR2 in each immunoprecipitated fraction was calculated using the 2^-ΔΔ*C*t^ method. First, the *C*t values for firefly luciferase in the input and in the anti-GFP immunoprecipitated fractions were subtracted from the corresponding *C*t values of HSUR1 or HSUR2 to normalize for variations in the number of cells (post-transfection) used to prepare the extracts and to account for differences among the samples in GFP-MS2 binding to the reporter transcripts and precipitation of the reporter mRNAs by the anti-GFP antibody.

Second, the relative abundance of wild-type HSUR1 or HSUR2 in each immunoprecipitated fraction was calculated by subtracting the normalized *C*t value for each HSUR in the input fraction from the corresponding normalized *C*t value in the anti-GFP immunoprecipitated fraction (Δ*C*t). The resulting value for wild-type HSUR1 or HSUR2 was then set as 1.0 (2^-Δ*C*t^_IP WT HSUR1/2_/2^-Δ*C*^t_IP WT HSUR1/2_). Third, the relative abundance of mutant HSUR1 or mutant HSUR2 in each immunoprecipitated fraction was calculated using the same procedure and expressed relative to the corresponding wild-type HSUR1 or HSUR2 (2^-ΔCt^_IP HSUR1/2 Mutant_/2^-Δ*C*t^_IP WT HSUR1/2_). Data were graphed with Prism 10 software. Statistical significance was determined separately for each reporter using ordinary one-way ANOVA, which indicated significant differences among the means for luciferase-JARID2_BS1 (*P* < 0.0001) and luciferase-STK4 (*P*< 0.0001), followed by an uncorrected Fisher’s least significant difference (LSD) multiple-comparisons test (Supplementary Table 2).

## RESULTS

### HSUR1 and HSUR2 bind Pumilio proteins

Comparison of publicly available HSUR1 and HSUR2 sequences from HVS and HVA (29–35) originally led to the identification of miRNA binding sites (4). Those analyses also identified a highly conserved motif within the 3′ loop of both HSUR1 and HSUR2 (Fig. 1A and Supplementary Fig. S1A and B). The RNA-Binding Protein Database (RBPDB) (36) identified these sequences as PREs. To determine whether HSUR1 and HSUR2 physically associate with PUM proteins, we performed immunoprecipitation experiments using antibodies against PUM1 and PUM2 in extracts prepared from wild-type HVS-A11-transformed marmoset T cells (cj319-WT), which express all seven HSURs encoded by HVS (4) (Fig. 1B). HSUR1 and HSUR2 were selectively enriched, although to different extents, in immunoprecipitates obtained with both PUM1 and PUM2 antibodies compared with control antibody immunoprecipitates, indicating that these two viral ncRNAs associate with both PUM1 and PUM2 in HVS-transformed cells.

Association of HSUR1 with miR-27 and engagement of ARE-BPs in HSUR1-mediated mRNA repression, as well as association of HSUR2 with miR-16, require interaction of these viral ncRNAs with miR-142-3p (7,9). To determine whether association of HSUR1 and HSUR2 with PUM1 and PUM2 similarly depends on interaction with miR-142-3p, we performed immunoprecipitation experiments using extracts from HeLa-H1 and HeLa-H2 cells (9) which constitutively express HSUR1 and HSUR2, respectively, but lack miR-142-3p. Extracts where prepared from these cells following transient transfection with either Control miRNA or miR-142-3p and subjected to immunoprecipitation with antibodies against PUM1 and PUM2 (Supplementary Fig. S1C). HSUR1 and HSUR2 showed similar association with PUM1 and PUM2 in the presence and absence of miR-142-3p, indicating that this miRNA is not required for association of either HSUR with PUM proteins.

We confirmed that the PREs present in HSUR1 and HSUR2 mediate the association of these viral ncRNAs with PUM proteins. Immunoprecipitation experiments with antibodies against PUM1 and PUM2 were performed on extracts prepared from HeLa cells that constitutively express mutant versions of HSUR1 and HSUR2 in which the PRE was mutagenized (H1PREm and H2PREm, Fig. 1A and C). Disruption of the PRE present in HSUR1 and HSUR2 did not affect their expression (see Input fractions, Fig. 1C) but abolished their ability to associate with PUM1 and PUM2 (Fig. 1C). These experiments confirm that HSUR1 and HSUR2 associate with PUM proteins through the PREs present in their 3′ loops.

### HSUR1 and HSUR2 do not alter PUM1 or PUM2 activity

PUM proteins are modulated by the long ncRNA *NORAD*, which acts as a competitive inhibitor by sequestering PUM proteins (23,24). Several characteristics of *NORAD* make it particularly well suited to inhibit PUM proteins:(i) both PUM proteins and *NORAD* are predominantly cytoplasmic; (ii) *NORAD* contains an unusually large number of PREs (∼17-20), enabling it to bind PUM proteins multivalently; and (iii) *NORAD* is relatively abundant compared with PUM proteins. In contrast, HSUR1 and HSUR2 contain only one PRE per molecule and predominantly localize to the nucleus. These properties argue against a model in which HSURs inhibit PUM proteins by sequestering them. However, HSURs do access the cytoplasm in at least two contexts: they have been shown to associate with mRNAs that are actively being translated (5,7), and they transiently enter the cytoplasm during their biogenesis, a process that is shared with cellular snRNAs (37). Thus, HSUR1 and HSUR2 could, in principle, bind PUM proteins in the cytoplasm and relocalize them to the nucleus, thereby altering their subcellular distribution and potentially affecting their function.

We performed subcellular fractionation experiments to determine whether HSUR1 and HSUR2 alter the localization of PUM proteins. Cytoplasmic and nuclear extracts were prepared from marmoset T cells transformed with either wild-type HVS (cj319-WT) (4), which expresses all HSURs, or a mutant version of HVS lacking HSUR1 and HSUR2 (cj319-Δ2a) (4,38), and analyzed them by Western and Northern blotting (Fig. 2A). No significant differences in the subcellular distribution of PUM1 and PUM2 were observed between the two cell lines, with both proteins being predominantly present in the cytoplasmic fraction. These results indicate that HSUR1 and HSUR2 do not affect the localization of PUM proteins.

**Figure 2.**
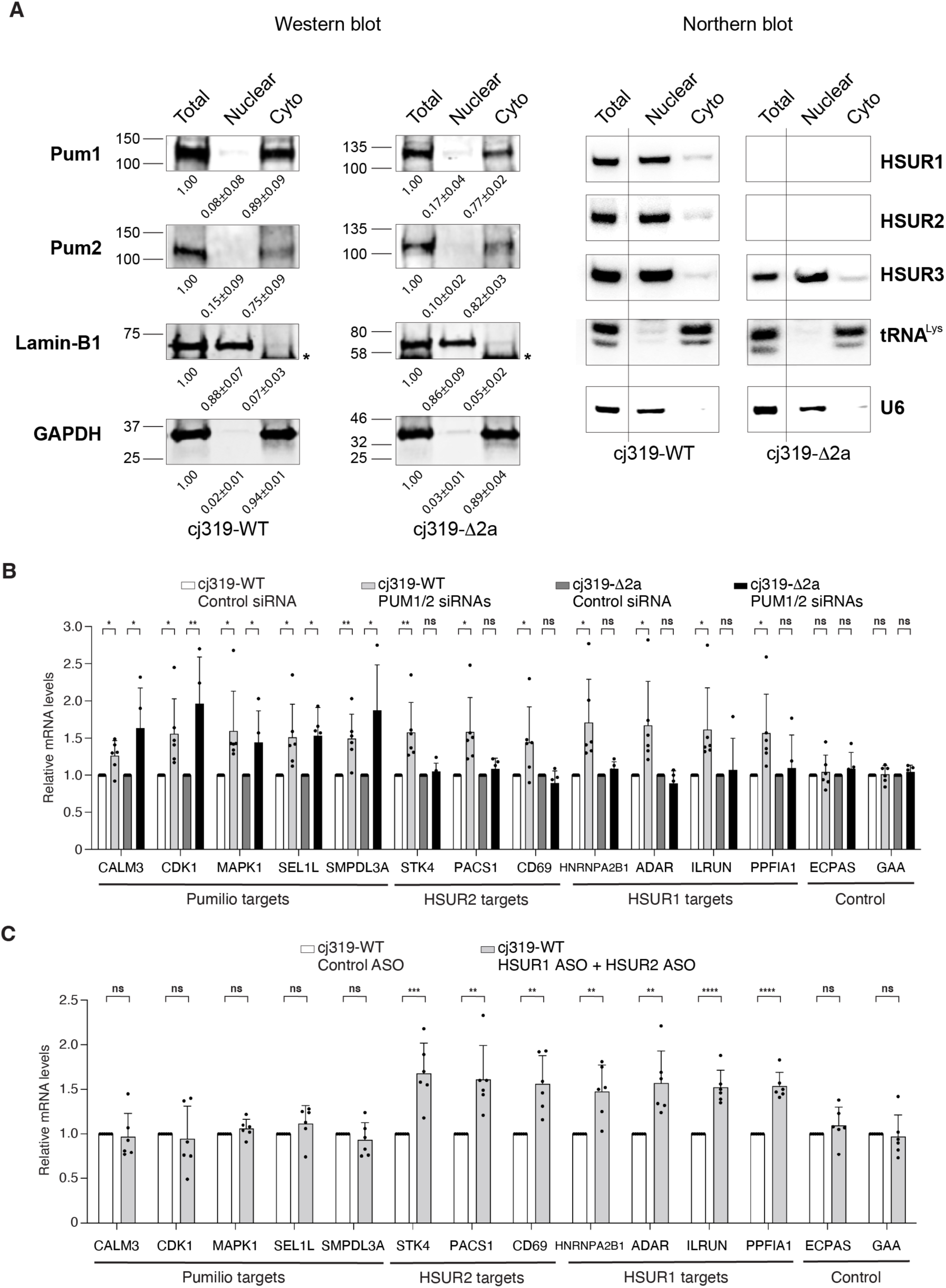
HSUR1 and HSUR2 do not alter PUM1 or PUM2 activity in HVS-infected cells. **(A)** Northern and Western blot analyses of cytoplasmic and nuclear extracts prepared from cj319-WT and cj319-Δ2a cells, showing the relative cytoplasmic and nuclear abundance of PUM proteins in the presence or absence of HSUR1 and HSUR2. Western blot analyses of the indicated proteins in nuclear and cytoplasmic (Cyto) fractions are shown on the left. Numbers below each lane represent the mean fractional amount of each protein present in the corresponding extract ± SD (*n* = 3). No statistically significant differences were observed in the nuclear and cytoplasmic distribution of PUM1 and PUM2 between cj319-WT and cj319-Δ2a cells (Supplementary Table 2). Membranes were re-probed for Lamin-B1 and GAPDH to verify successful subcellular fractionation. The asterisk denotes a non-specific band detected by the secondary antibody. Northern blot analysis of the same fractions is shown on the right. Membranes were probed for the indicated RNAs. U6 snRNA and lysine tRNA were used to verify successful subcellular fractionation. Vertical lines indicate that the lanes containing the total extracts from cj319-WT and cj319-Δ2a cells were not contiguous with the other lanes shown. **(B)** qRT-PCR analysis of PUM proteins and HSUR1 and HSUR2 target mRNAs in cj319-WT and cj319-Δ2a cells transiently transfected with Control siRNA or siRNAs targeting PUM1 and PUM2. ECPAS and GAA served as control mRNAs. **(C)** Same as in (B), using cj319-WT cells transiently transfected with Control ASO or ASOs targeting HSUR1 and HSUR2. \**P* < 0.05; \*\**P* < 0.01; \*\*\**P* < 0.001; \*\*\*\**P* < 0.0001.

We next tested whether HSUR1 and HSUR2 could affect the ability of PUM proteins to repress their target mRNAs. We reasoned that if HSUR1 and HSUR2 inhibit PUM proteins and render them unable to regulate their target mRNAs, depletion of PUM proteins in a cell line expressing HSUR1 and HSUR2 should have little or no effect on the abundance of PUM target mRNAs. In contrast, depletion of PUM proteins in a cell line lacking HSUR1 and HSUR2 should result in upregulation of PUM target mRNAs. To test this hypothesis, we used siRNAs to deplete both PUM1 and PUM2 in cj319-WT cells, which express HSUR1 and HSUR2, and in cj319-Δ2a cells, which lack HSUR1 and HSUR2 (Supplementary Fig. 2A), and analyzed the abundance of PUM target mRNAs by qRT-PCR (Fig. 2B). We evaluated PUM targets that had been previously validated experimentally by depleting both PUMs from human cells and measuring differential RNA expression (18). We selected specific PUM target mRNAs based on the following criteria: i) they contain PREs in their 3′UTRs; ii) they showed the greatest sensitivity to PUM protein levels in human cells (18); and iii) they do not associate with HSUR1 and HSUR2 (6,7) and are therefore not expected to be directly regulated by these two viral ncRNAs. Depletion of PUM1 and PUM2 in cj319-Δ2a cells, which lack HSUR1 and HSUR2 (Fig. 2A), resulted in upregulation of all PUM target mRNAs tested (Fig. 2B), indicating that these mRNAs are directly regulated by PUM proteins in HVS-transformed marmoset T cells, as previously demonstrated in human cells (18). PUM target mRNAs were similarly upregulated in cj319-WT cells, which express HSUR1 and HSUR2 (Fig. 2A), following depletion of PUM proteins, indicating that PUM proteins actively regulate their target mRNAs even in the presence of HSUR1 and HSUR2. These results do not support a model in which HSUR1 and HSUR2 impair PUM protein function. As controls, we analyzed ECPAS and GAA, which do not contain PREs and are not direct targets of HSUR1 or HSUR2, and observed no changes in their abundance in either cell line (Fig. 2B). We next evaluated experimentally validated HSUR1 and HSUR2 target mRNAs (6,7) that do not contain PREs and are therefore not expected to be directly regulated by PUM proteins. Surprisingly, depletion of PUM proteins resulted in upregulation of these HSUR1 and HSUR2 target mRNAs in cj319-WT cells, but not in cj319-Δ2a cells, which lack HSUR1 and HSUR2 (Fig. 2B). These results indicate that PUM proteins can indirectly affect the expression of HSUR target mRNAs in a HSUR-dependent manner. Altogether, these results argue against a model in which HSURs inhibit PUM protein function and instead support a model in which PUM proteins are required for HSUR-mediated mRNA repression.

The results described above indicate that PUM proteins actively regulate their target mRNAs in the presence or absence of HSUR1 and HSUR2. However, the extent of derepression observed following PUM protein depletion differed slightly for some of the targets between the two cell lines. One caveat of utilizing cj319-WT and cj319-Δ2a cells is that their transcriptomes are expected to diverge considerably, because both HSUR1 and HSUR2 affect gene expression by binding to, and regulating target mRNAs (5–7), with HSUR1 alone responsible for altering the abundance of ∼37% of mRNAs in infected cells by ≥1.5-fold (7).

Differences in the transcriptomes of the two cell lines could therefore explain the modest differences in the extent of derepression observed for some PUM target mRNAs. In addition, these two cell lines have been independently propagated for over two decades (39), potentially allowing the independent accumulation of mutations that could also contribute to the differences observed in the experiments described above. To confirm that HSUR1 and HSUR2 do not affect the activity of PUM proteins in infected cells, we depleted both HSUR1 and HSUR2 with antisense oligonucleotides (ASOs) which had previously been shown to efficiently knockdown expression of these two viral ncRNAs following transient transfection into HVS-transformed marmoset T cells (5,7). Efficient co-depletion of HSUR1 and HSUR2 from cj319-WT cells (Supplementary Fig. 2B) resulted in increased abundance of HSUR1 and HSUR2 target mRNAs (Fig. 2C), whereas no changes were observed in any of the PUM target mRNAs tested or in control mRNAs. Thus, PUM-dependent regulation of PUM target mRNAs was maintained despite the presence of HSUR1 and HSUR2, and depletion of HSUR1 and HSUR2 did not measurably alter the expression of these PUM target mRNAs. Together, these complementary experiments provide evidence that HSUR1 and HSUR2 do not inhibit or otherwise measurably alter PUM protein activity in HVS-infected cells. Instead, the data support a model in which HSUR1 and HSUR2 require the presence of PUM proteins to regulate their target mRNAs.

### Pumilio proteins are required for HSUR-mediated mRNA repression

To determine whether PUM proteins are required for HSUR1- and HSUR2-mediated mRNA repression, we evaluated the ability of H1PREm and H2PREm mutants, which get expressed in HeLa cells at similar levels to those of wild-type HSUR1 and HSUR2 but cannot bind PUM proteins (Fig. 1C), to repress HSUR1 and HSUR2 target mRNAs, respectively. Transient expression of miR-142-3p in HeLa cells expressing wild-type HSUR1 (HeLa-H1) and HSUR2 (HeLa-H2) resulted in repression of previously validated HSUR1 and HSUR2 target mRNAs (7,9) whereas no changes were observed in control mRNAs (Fig. 3A and B). In contrast, mutant versions of HSUR1 and HSUR2 that cannot bind PUM proteins failed to significantly affect the expression of HSUR1 and HSUR2 target mRNAs in the presence of miR-142-3p (Fig. 3A and B). Because both HSUR1 and HSUR2 require base-pairing with miR-142-3p to repress their target mRNAs (7,9), we next considered whether disruption of PUM protein interaction could affect the ability of HSUR1 and HSUR2 to bind miR-142-3p and, consequently, their ability to repress target mRNAs. We monitored HSUR association with miR-142-3p by immunoprecipitation with antibodies against Sm proteins, which recognize HSURs (2) and selectively immunoprecipitate miR-142-3p only in the presence of HSUR1 or HSUR2 (4,9), followed by Northern blotting (Supplementary Fig. 3). Mutant versions of HSUR1 and HSUR2 that cannot bind PUM proteins were precipitated with Sm antibodies to the same extent as wild-type HSURs, indicating that disruption of PUM protein binding does not interfere with Sm protein binding or HSUR RNP formation. Similarly, we observed no differences in the ability of wild-type HSURs and their mutant counterparts to associate with miR-142-3p, indicating that the inability of H1PREm and H2PREm to repress target mRNAs is not a consequence of impaired association with miR-142-3p.

**Figure 3.**
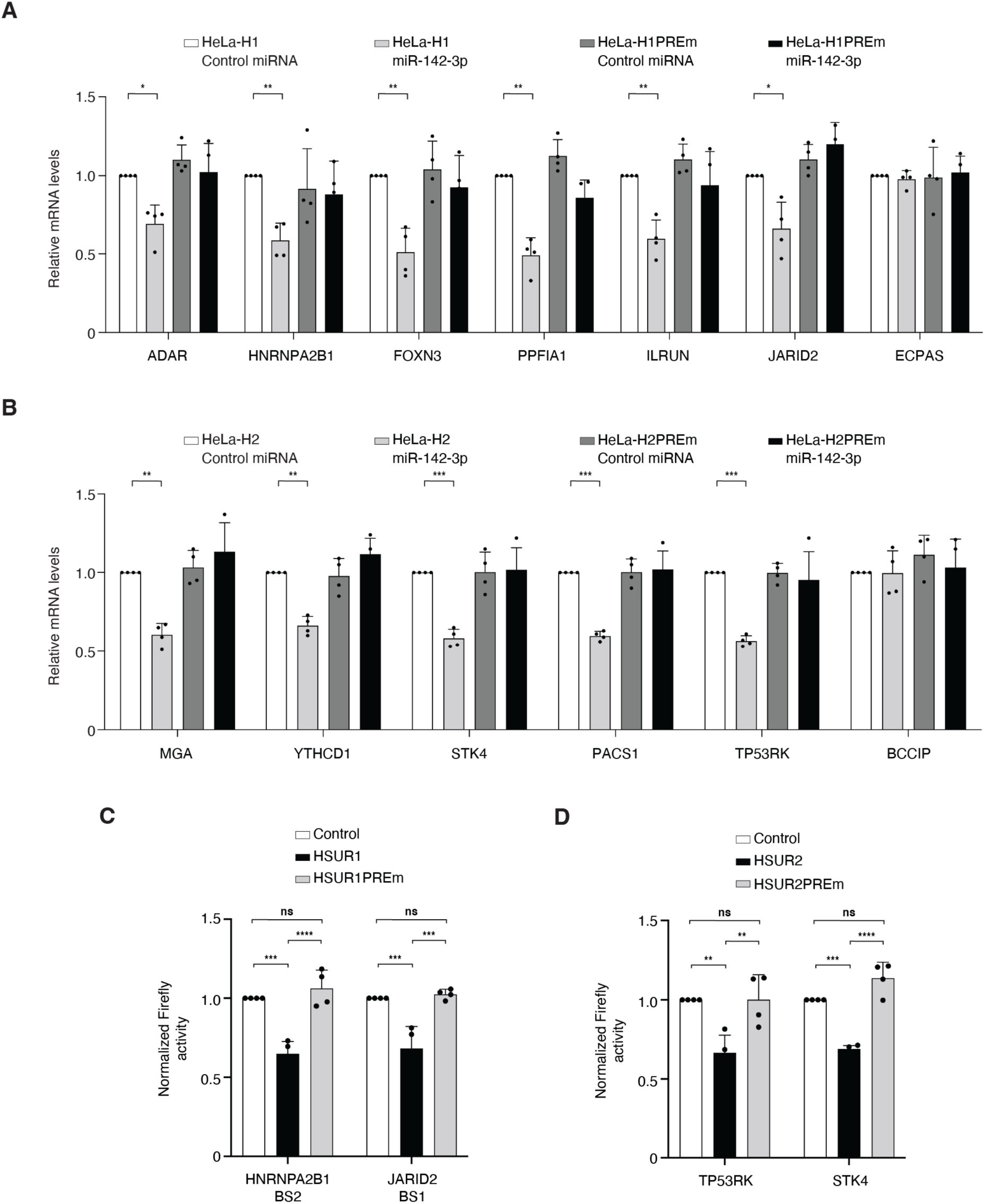
PUM proteins are required for HSUR-mediated mRNA repression. **(A-B)** qRT-PCR analyses of HSUR1 (A) or HSUR2 (B) target mRNAs in HeLa cells stably expressing wild-type HSUR1 (HeLa-H1) or a mutant version in which the PRE sequence was disrupted (HeLa-H1PREm) (A), or wild-type HSUR2 (HeLa-H2) or a mutant version in which the PRE sequence was disrupted (HeLa-H2PREm) (B), following transfection with either Control miRNA or miR-142-3p. **(C)** Luciferase assays in HuT78 cells stably expressing reporter transcripts containing 3′UTR fragments with single HSUR1-binding sites, following transient transfection with plasmids expressing wild-type HSUR1 or the PRE-mutant H1PREm. Firefly luciferase levels were normalized to *Renilla* luciferase levels. **(D)** Same as in (C), using U937 cells expressing reporter transcripts containing 3′UTR fragments with single HSUR2-binding sites transiently transfected with plasmids expressing wild-type HSUR2 or the mutant H2PREm. \**P* < 0.05; \*\**P* < 0.01; \*\*\**P* < 0.001; \*\*\*\**P* < 0.0001. When not indicated, differences were not statistically significant.

We next tested the ability of wild-type and PRE mutant versions of HSUR1 and HSUR2 to repress luciferase reporter transcripts carrying a single binding site for either viral ncRNA. These reporter transcripts provide two advantages: interactions between each HSUR and the individual binding sites have been previously validated experimentally by mutational analyses (6,7), which demonstrated that base-pairing between the HSUR and its target site is required for repression; and the reporter transcripts do not contain PREs and therefore should not be sensitive to changes in PUM protein activity independently of HSURs. We transiently expressed HSUR1, HSUR2, H1PREm, or H2PREm in either U937 or HuT78 human T cells, which naturally express miR-142-3p and constitutively express luciferase reporter genes containing partial 3′UTR sequences with single binding sites for HSUR1 (HNRNPA2B1 BS2 and JARID2 BS1 in HuT78 cells; Fig. 3C) or HSUR2 (TP53RK or STK4 in U937 cells; Fig. 3D). These experiments recapitulated the effects observed with the endogenous target mRNAs.

Disruption of the PRE sequences present in HSUR1 and HSUR2 impairs association with PUM proteins (Fig. 1C) and render these viral ncRNAs unable to repress their target mRNAs (Fig. 3), suggesting that association of HSURs with PUM proteins is required for HSUR-mediated mRNA repression. However, an alternative hypothesis is that the sequences within HSURs that constitute the PREs have a second function, independent of PUM protein binding, that is essential for HSUR-mediated mRNA repression. To distinguish between these possibilities, we generated versions of HSUR1 and HSUR2 in which the upper portion of their 3′ stem and loop were replaced with the bacteriophage λ boxB hairpin (HSUR1*boxB* and HSUR2*boxB*; Fig. 4A). For assembly of Sm proteins onto snRNAs, the specific sequence of the stem-loop adjacent to the Sm binding site is not important, but it must be located within a short distance of the 3′ end of the RNA(40). Thus, replacing the upper portion of the stem and loop with the boxB hairpin does not impair HSUR expression (Supplementary Fig. 4A) because it does not alter the distance between the Sm binding sites, stem-loop and the 3′ end of the RNAs (Fig. 4A). We also generated plasmids for transient expression of PUM1 or PUM2 fused to the short N-terminal domain (amino acids 1-22) of the bacteriophage λ transcriptional antiterminator protein N (λN-PUM1 and λN-PUM2, Fig. 4A). The λN domain binds to the boxB hairpin RNA (41), thereby providing a means to tether PUM1 or PUM2 directly to the modified boxB-carrying HSURs that no longer contain a PRE. Transient expression of HSUR1*boxB* or HSUR2*boxB* alone did not result in repression of luciferase reporter transcripts carrying the corresponding binding sites (Fig. 4B and C, and Supplementary Fig. 4F and G). In contrast, repression of the same reporter transcripts was restored when either λN-PUM1 and λN-PUM2 was co-expressed with HSUR1*boxB* or HSUR2*boxB* (Fig. 4B and C, and Supplementary Fig. 4F, and G), whereas λN-PUM1 and λN-PUM2 alone did not affect expression of these reporter transcripts (Supplementary Fig. 4B-E). These results indicate that the primary function of the PRE in HSUR1 and HSUR2 is to recruit PUM proteins and that PUM binding, rather than another function encoded by the PRE sequence itself, is sufficient to enable these viral ncRNAs to repress target mRNAs.

**Figure 4.**
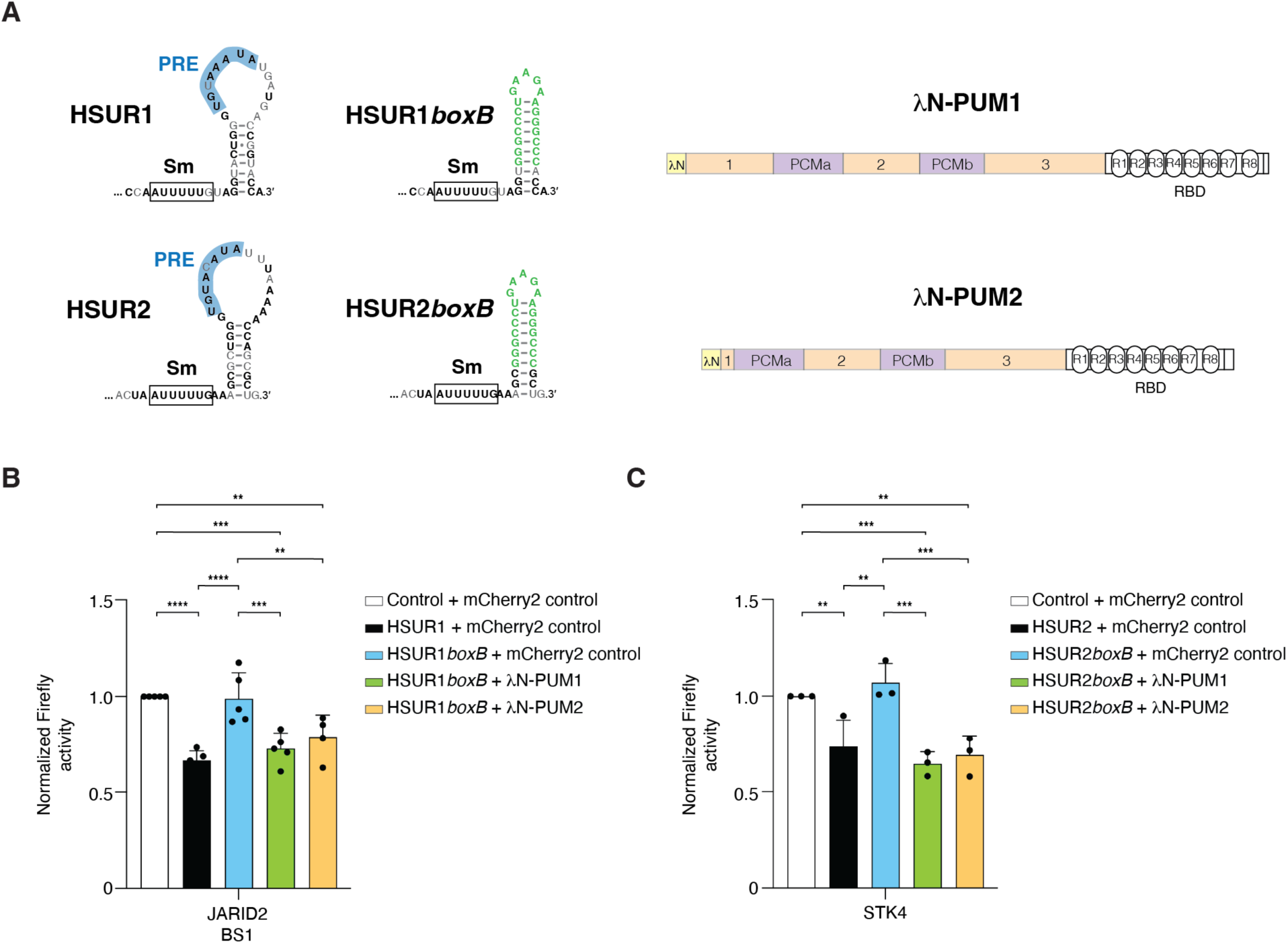
Association of PUM proteins with HSURs is required for HSUR-mediated mRNA repression. **(A)** Reagents used to test whether recruitment of PUM proteins to HSURs is required for HSUR-mediated mRNA repression. Nucleotide sequence of the 3′ portions of HSUR1 and HSUR2 and mutant versions in which the 3′ stem-loop was partially replaced with boxB hairpins (HSUR1*boxB* and HSUR2*boxB*) are shown. Sequences constituting PREs are highlighted in blue, and modified nucleotides constituting the boxB hairpins are shown in green. Diagrams of PUM1 and PUM2 fused to λN are shown on the right. The λN domain binds the boxB hairpin, thereby tethering PUM1 or PUM2 to HSUR1*boxB* or HSUR2*boxB.* Numbers 1-3 indicate repression domains. Pumilio Conserved Motifs (PCMa and PCMb), as well as the Pum repeats (R1-R8) constituting the RNA Binding Domain (RBD), are indicated. **(B)** Luciferase assays in Hut78 cells expressing a reporter transcript containing a fragment of JARID2 3′UTR with a single HSUR1 binding site (binding site 1, BS1), following transient co-transfection of a control plasmid expressing mCherry2 alone (mCherry2 control) or plasmids encoding mCherry2 and λN-PUM1 or λN-PUM2, together with a control plasmid containing the U1 promoter (Control) or a plasmid expressing wild-type HSUR1 or HSUR1*boxB* under the control of U1 promoter. Firefly luciferase levels were normalized to *Renilla* luciferase levels. **(C)** Same as in (B), using U937 cells expressing a reporter transcript containing a fragment of the STK4 3′UTR with a single HSUR2 binding site and a control plasmid containing HSUR2 promoter or plasmids expressing wild-type HSUR2 or HSUR2*boxB* under the control of the HSUR2 promoter. \*\**P* < 0.01; \*\*\**P* < 0.001; \*\*\*\**P* < 0.0001. When not indicated, differences were not statistically significant.

### Pumilio proteins enable HSUR association with target mRNAs

To repress target mRNAs, HSUR1 and HSUR2 must interact through base-pairing with both miR-142-3p and their respective target mRNAs (5–7,9). Disruption of the interactions between HSURs and PUM proteins renders HSUR1 and HSUR2 unable to repress target mRNAs (Fig. 3) but does not impair their association with miR-142-3p (Supplementary Fig. 3). We therefore next tested whether versions of HSUR1 and HSUR2 that cannot bind PUM proteins can still associate with target mRNAs. The luciferase reporter genes carrying a binding site for HSUR1 (JARID2 BS1) or HSUR2 (STK4) were modified by introducing MS2 RNA hairpins between the HSUR binding sites and the cleavage and polyadenylation signal (Fig. 5A). Introduction of the MS2 hairpins did not affect the sensitivity of these reporter transcripts to HSUR1 and HSUR2 expression (Supplementary Fig. 5A and B). HuT78 cells, which endogenously express miR-142-3p and constitutively express the luciferase reporter genes with MS2 hairpins, were transiently co-transfected with a plasmid expressing MS2 coat protein fused to eGFP (28) and a plasmid expressing wild-type HSUR1 or HSUR2, or mutant versions of these viral ncRNAs that cannot bind miR-142-3p, cannot base-pair with binding sites present in the luciferase reporter transcripts (6,7), or cannot associate with PUM proteins (Fig. 5A and Supplementary Fig. 5C and D). Following transfection, we performed immunoprecipitation with antibodies against GFP to isolate the reporter transcripts through the MS2-eGFP fusion protein. The presence of HSUR1, HSUR2, and their mutant variants in the immunoprecipitates was then monitored by qRT-PCR using internal primers that do not overlap with any of the mutagenized residues in each viral ncRNA (Supplementary Fig. 5C and D). As expected, versions of HSUR1 and HSUR2 carrying mutations previously shown to disrupt base-pairing with JARID2 and STK4 binding sites, respectively (Supplementary Fig. 5C and D), were not enriched in GFP immunoprecipitates, indicating that they do not associate with the reporter transcripts. Similarly, H1PREm and H2PREm, which cannot bind PUM proteins, were not enriched in GFP immunoprecipitates, indicating that they do not associate with luciferase reporter transcripts despite retaining an intact mRNA binding region. In contrast, we found no significant difference in the ability of mutant versions of HSURs that cannot bind miR-142-3p to associate with target mRNAs when compared with wild-type HSURs (Fig. 5C and D). These results are consistent with previous results showing that HSUR2 can be psoralen-crosslinked to mRNAs in the absence of miR-142-3p (5). The observation that mutant HSURs unable to bind miR-142-3p cannot repress target mRNAs despite retaining the ability to bind PUM proteins (Supplementary Fig. 1C) and target mRNAs further indicates that PUM proteins do not function canonically by recruiting deadenylation complexes to directly mediate HSUR-dependent mRNA repression.

**Figure 5.**
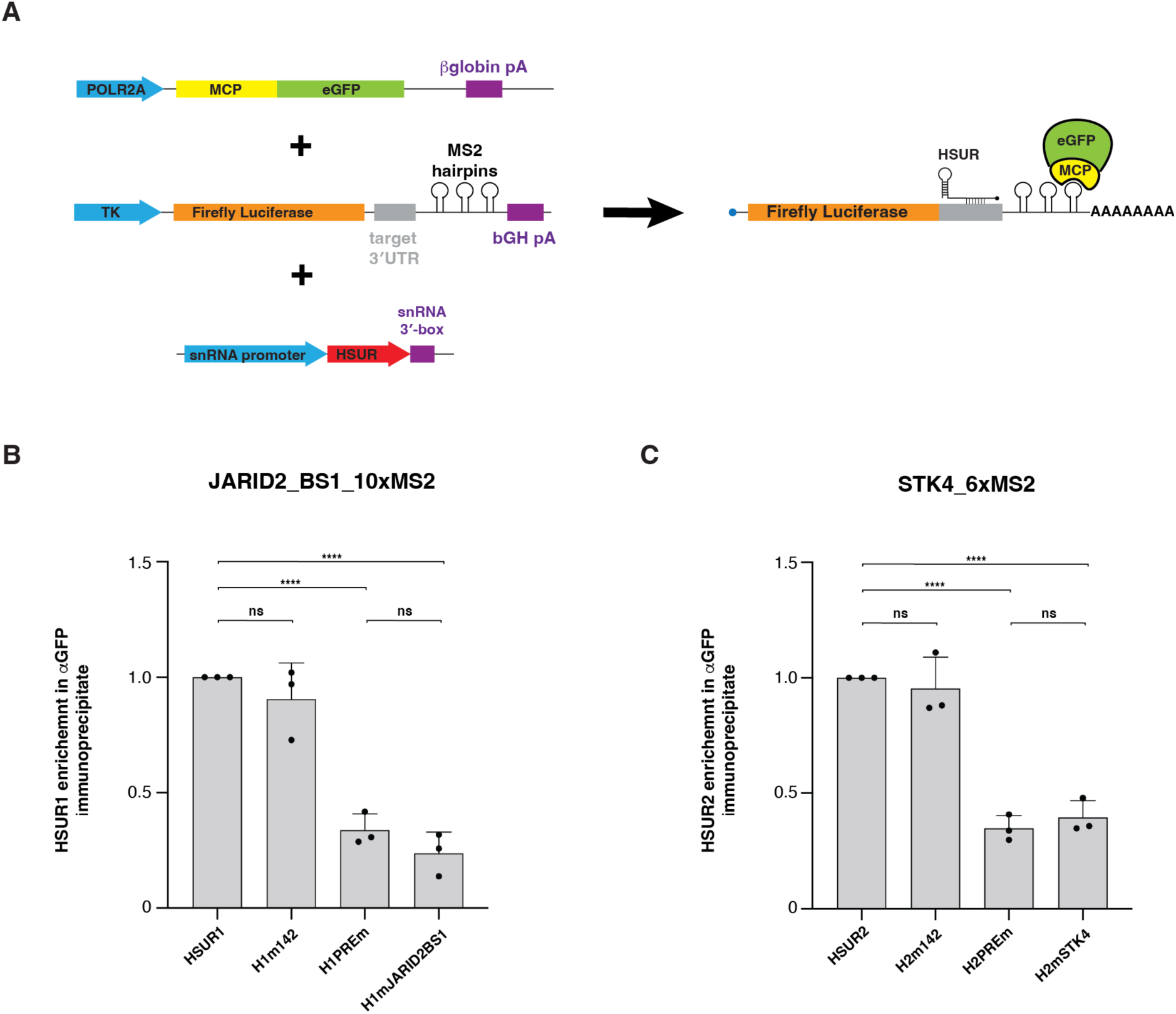
PUM proteins license HSUR association with target mRNAs. **(A)** Schematic of strategy used to determine whether PUM proteins are required for HSUR association with target mRNAs. HuT78 cells stably expressing luciferase reporter transcripts containing fragments of 3′UTRs with single HSUR binding sites and MS2 hairpins were transiently co-transfected with plasmids expressing MS2 coat protein (MCP) fused to eGFP and either wild-type HSUR1 of HSUR2 or mutant versions of these HSURs. For HSUR1, the mutants included H1m142, which cannot bind miR-142-3p; H1PREm, which cannot bind PUM proteins; and H1mJARID2BS1, which carries mutations that impair base-pairing with the binding site within the JARID2 reporter. For HSUR2, the corresponding mutants included H2m142, which cannot bind miR-142-3p; H2PREm, which cannot bind PUM proteins; and H2mSTK4, which carries mutations mutation that impair base-pairing with the binding site within the STK4 reporter. Following transfection, cell extracts were prepared, and HSUR-reporter mRNA-MCP/eGFP complexes were isolated by immunoprecipitation with antibodies against GFP and analyzed by qRT-PCR to determine HSUR enrichment. **(B)** Enrichment of wild-type HSUR1 or HSUR2 and the indicated mutant version of these viral ncRNAs in immunoprecipitates obtained with antibodies against GFP. \*\*\*\**P* < 0.0001.

Together, these results support a model in which PUM proteins bind the PRE within HSUR1 and HSUR2 and thereby enable these viral ncRNAs to engage their target mRNAs through sequence-specific base-pairing. Thus, rather than functioning as downstream effectors of HSUR-mediated repression, PUM proteins act as factors that facilitate HSUR association with target mRNAs, which is required for subsequent mRNA repression.

## DISCUSSION

NcRNAs regulate gene expression through diverse mechanisms that frequently depend on their ability to interact with RBPs. In many characterized pathways, however, RBPs function primarily as downstream effectors that execute regulatory instructions encoded by the ncRNA. For example, miRNAs guide Argonaute proteins to target mRNAs, after which Argonaute recruits additional factors that promote transcript destabilization and translational repression (42). In this pathway, the miRNA itself is dispensable for repression, since artificially tethering Argonaute to mRNAs is sufficient to exert repression (43). Our findings reveal a fundamentally different role for an RBP in ncRNA-mediated regulation. We show that PUM1 and PUM2 are required for HSUR1 and HSUR2 to associate with target mRNAs and thereby repress gene expression (Fig. 4). Rather than functioning as downstream effectors of HSUR-mediated repression, PUM proteins bind PRE motifs within these viral ncRNAs and enable their association with target mRNAs (Fig. 5). Thus, in this context, rather than executing the ncRNA regulatory activity, PUM proteins function as RBPs that facilitate target recognition by ncRNAs.

The mechanism uncovered here is distinct from the canonical activity of PUM proteins. PUM1 and PUM2 recognize PREs within target mRNAs and can promote transcript repression through recruitment of the CCR4-NOT deadenylase complex (17,44). Our depletion experiments demonstrate that PUM proteins retain their ability to regulate their canonical target mRNAs in cells expressing HSUR1 and HSUR2 (Fig. 2B). Conversely, depletion of HSUR1 and HSUR2 does not measurably alter the expression of PUM target mRNAs (Fig. 2C). These complementary observations argue against a model in which HSURs globally inhibit or otherwise substantially alter PUM protein activity. Instead, HSURs use PUM proteins independently of their canonical role in recognizing and repressing PUM target mRNAs. This finding expands the functional repertoire of PUM proteins and illustrates how an RNA-binding protein can be repurposed by a viral ncRNA to perform a function distinct from its established activity.

HSUR1 and HSUR2 present an unusual challenge for target recognition. Unlike miRNAs, which contain a relatively short seed sequence that provides a common determinant for target recognition, HSUR1 and HSUR2 base-pair with their target mRNAs through variable regions with no fixed position or sequence (6,7). As a result, individual HSUR target sites can differ substantially in sequence configuration, and neither HSUR contains a single defined seed region that can account for recognition of all of its targets. Our results uncover another piece of information that provide a glimpse into this puzzle. PUM1 and PUM2 recognize a conserved PRE located within HSUR1 and HSUR2 themselves (Fig. 1), and disrupting this PRE prevents the viral ncRNAs from associating with target mRNAs (Fig. 5). Importantly, disruption of PUM binding does not affect HSUR association with miR-142-3p (Supplementary Fig. 3), indicating that the requirement for PUM proteins is not explained by impaired formation of the HSUR-miRNA complex. Furthermore, tethering of PUM1 or PUM2 directly to HSUR1 or HSUR2 through the λN–boxB system restores repression in the absence of the native PRE (Fig. 4).

Together, these findings indicate that PUM recruitment is not merely correlated with HSUR function but is itself the critical event that enables HSURs to engage their target mRNAs.

The remarkable flexibility of HSURs in sequence recognition raises the question of why they do not bind a much larger number of target mRNAs. The requirement for PUM proteins introduces an additional layer of specificity into HSUR-mediated regulation that may explain this selectivity. HSUR1 and HSUR2 contain sequence information that determines their association with miR-142-3p, as well as with miR-16 in the case of HSUR2 and ARE-BPs in the case of HSUR1, and their target mRNAs. However, productive interaction with target mRNAs additionally requires recruitment of PUM proteins. Thus, HSUR-mediated target recognition may involve two distinct requirements: recognition of the target mRNA through HSUR base-pairing and PUM-dependent selection or stabilization of productive interactions within the appropriate ribonucleoprotein context. PUM proteins could, for example, interact with proteins associated with target mRNAs, thereby selecting the appropriate mRNP context as an initial step in target recognition. Alternatively, PUM proteins could directly stabilize the interaction between HSURs and their target RNAs after base-pairing is established or induce a conformation of the HSUR that is competent for target recognition. In either case, the requirement for PUM proteins could prevent the extensive promiscuity that might otherwise result from the broad sequence flexibility of HSUR base-pairing. The viral ncRNA determines the identity of the target RNA through sequence complementarity, while PUM binding provides an additional requirement for productive engagement. The combination of these interactions could allow HSURs to discriminate between transient or nonproductive RNA contacts and interactions that are competent to initiate repression. The precise molecular mechanism by which PUM binding promotes HSUR association with target mRNAs remains to be determined. Our experiments establish the requirement for PUM, but further experimentation will be required to distinguish among these possibilities.

An important implication of this mechanism is that PUM proteins may provide a general model that allow ncRNAs lacking canonical target-recognition elements to engage with target transcripts. PUM proteins are generally considered sequence-specific regulators because their canonical function depends on recognizing PREs within target mRNAs. Our findings instead demonstrate that the PRE can reside within an ncRNA whose function is to regulate a different set of RNAs. In this configuration, PUM recognition is effectively transferred from the target mRNA to the regulatory ncRNA. This raises the possibility that other cellular or viral ncRNAs containing PUM-binding elements may similarly exploit PUM proteins to regulate RNA-RNA interactions. Whether such mechanisms occur broadly remains to be determined, but the existence of this pathway suggests that the repertoire of PUM-dependent RNA regulation may extend beyond direct recognition of PRE-containing mRNAs.

Several questions remain regarding how PUM binding mechanistically promotes HSUR-mRNA association. Structural and biochemical studies will be required to determine whether PUM binding alters the conformation of HSUR1 or HSUR2, directly contacts target RNAs, or recruits additional RNA-binding proteins that stabilize HSUR–mRNA interactions. It will also be important to determine whether PUM proteins remain associated with HSURs after target mRNA engagement and whether their canonical interactions with CCR4-NOT or other effector proteins contribute at any stage of HSUR-mediated repression. Our results indicate that canonical PUM-dependent repression is not required for HSUR function, but they do not exclude additional roles for PUM-associated proteins after HSUR recruitment to target transcripts.

The shared structural and mechanistic characteristics of HSUR1 and HSUR2 also raise the possibility that these two ncRNAs originated through an ancestral gene duplication event followed by functional diversification (45,46). HSUR1 and HSUR2 occupy adjacent positions in the HVS and HVA genomes (29,30) and retain related sequence and structural features, while also sharing several mechanistic properties, including their requirement for association with miR-142-3p for mRNA repression and with PUM proteins for target recognition. A duplication and divergence model could therefore explain how two related ancestral ncRNAs acquired distinct target specificities and regulatory functions while retaining a common miR-142-3p-dependent mechanism for mRNA repression and PUM-dependent mechanism for engaging target mRNAs. Under this model, conservation of the PREs would reflect an ancestral function retained by both RNAs, whereas divergence of other regions could have allowed HSUR1 and HSUR2 to evolve distinct miRNA interactions and target repertoires. Although the available sequence conservation does not by itself establish that HSUR1 and HSUR2 arose by duplication, their genomic proximity, sequence relatedness, and shared mechanistic dependence on miR-142-3p and PUM proteins are consistent with such an evolutionary origin.

In summary, our findings identify a non-canonical function for PUM1 and PUM2 in which these proteins act as enabling factors for ncRNA-mediated target recognition. By binding a PRE within HSUR1 or HSUR2, PUM proteins allow these viral ncRNAs to associate with target mRNAs through sequence-specific base-pairing and thereby promote repression. This mechanism differs from both the canonical function of PUM proteins on PRE-containing mRNAs and the competitive sequestration of PUM proteins by NORAD. More broadly, these findings demonstrate that RNA-binding proteins traditionally viewed as direct regulators of mRNAs can instead function as molecular adaptors that enable ncRNAs to engage their targets, expanding the mechanistic strategies through which ncRNAs control gene expression.

## Acknowledgements

We thank the members of the Flow Cytometry Core Facility at the University of Utah for help with cell sorting.

## Author contributions

D.C. conceived the project, supervised the project and obtained funding. C.G. and D.C. designed and performed all experiments and performed data interpretation. D.C. and C.G. wrote the manuscript, discussed the results and revised the manuscript.

## Declaration of generative AI and AI-assisted technologies in the writing process

During the preparation of this work the authors used ChatGPT-5 to improve readability of the manuscript. After using this tool/service, the authors reviewed and edited the content as needed and take full responsibility for the content of the published article.

---

**Supplementary Figure 1.**
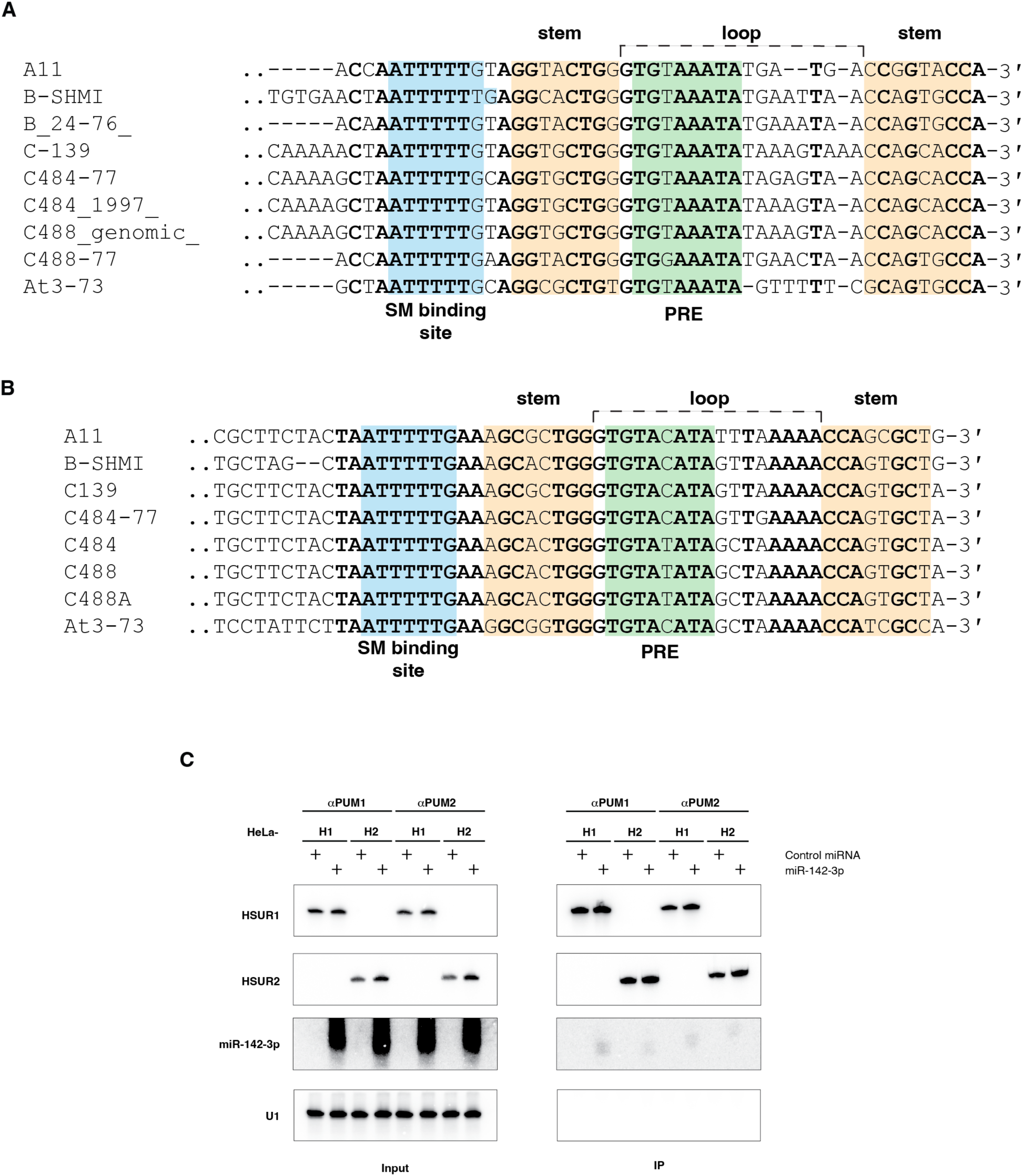
HSUR1 and HSUR2 associate with Pumilio proteins. **(A)** Alignment of reported genomic sequences (see main text) spanning the SM binding site to the 3′ end of HSUR1. The Sm binding site is highlighted in blue, sequences corresponding to stems are highlighted in orange, and sequences corresponding to the Pumilio Response Element (PRE) are highlighted in green. Perfectly conserved residues are shown in bold. **(B)** Same as in (A) for HSUR2. **(C)** Co-immunoprecipitation of HSUR1 and HSUR2 with antibodies against PUM1 or PUM2 from extracts prepared from HeLa cells expressing either HSUR1 (HeLa-H1) or HSUR2 (HeLa-H2) following transfection with either a control miRNA or miR-142-3p. Northern blotting was used to detect HSUR1, HSUR2, miR-142-3p, and cellular U1 snRNA as control for immunoprecipitation and sample loading. Input, 5%; IP, 95%.

**Supplementary Figure 2.**
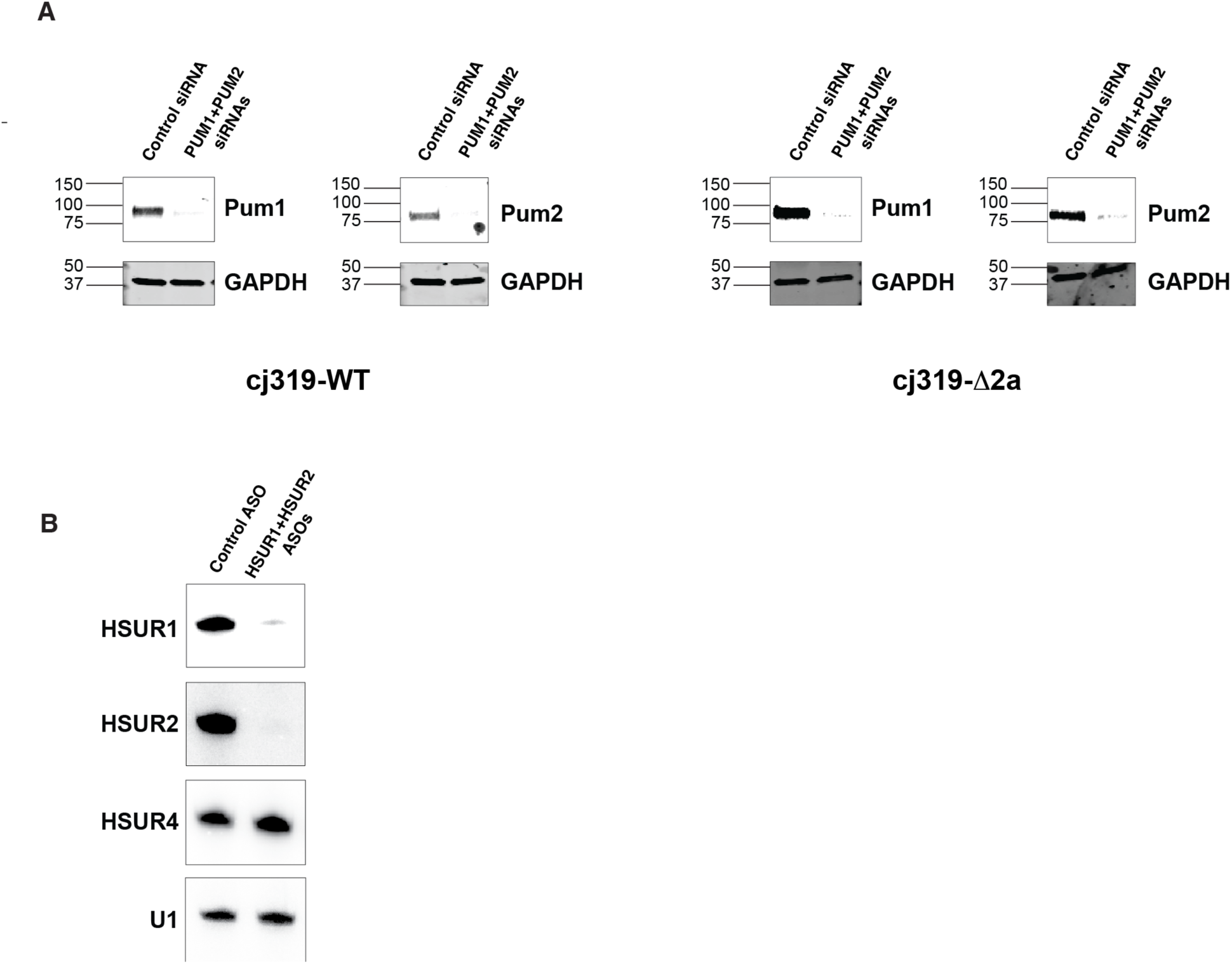
Knockdown of PUM proteins and HSUR expression in HVS-infected cells. **(A)** Western blot analyses of PUM1 and PUM2 in extracts from cj319-WT and cj319-Δ2a cells transiently transfected with either a control siRNA or siRNAs targeting PUM1 and PUM2. GAPDH serves as a loading control. (**B)** Northern blot analysis of HSUR1, HSUR2, HSUR4, and cellular U1 snRNAs in total RNA from cj319-WT cells transiently transfected with either a control ASO or ASOs targeting HSUR1 and HSUR2.

**Supplementary Figure 3.**
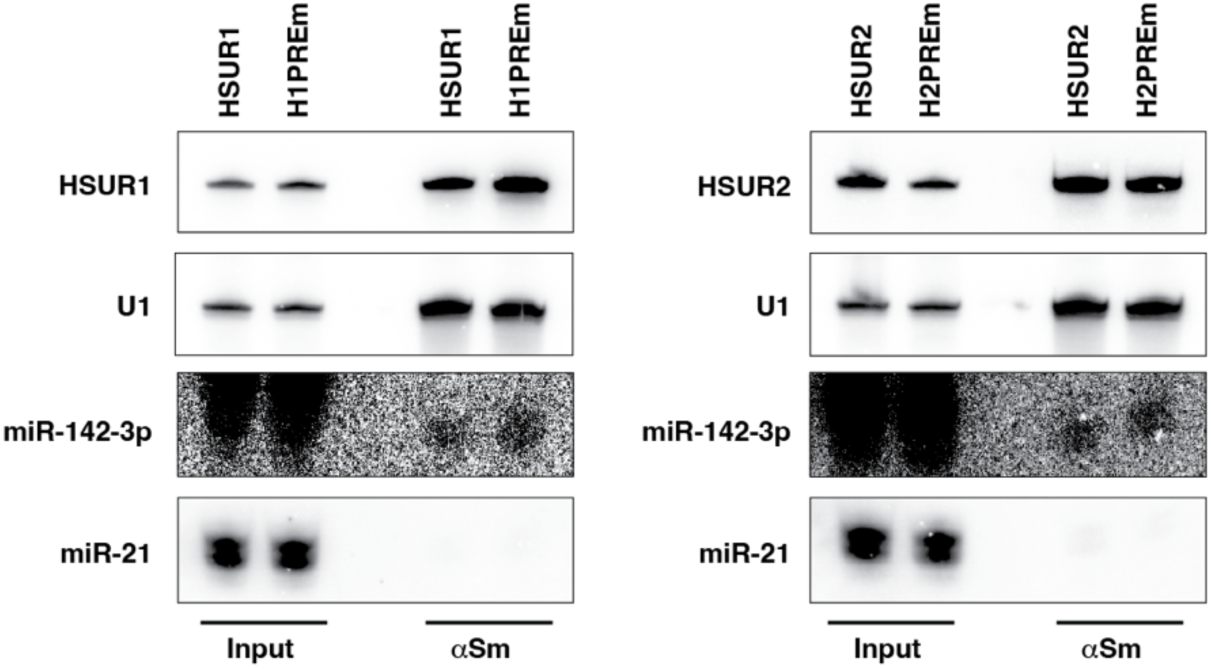
PUM proteins are not required for HSUR-miR-142-3p association. Co-immunoprecipitation of miRNAs with αSm antibodies from HeLa-H1, HeLa-H1PREm, HeLa-H2, and HeLa-H2PREm cells following transfection with miR-142-3p. Northern blot was probed for HSUR1, HSUR2, miR-142-3p and U1 snRNA as immunoprecipitation control. Input, 5%; IP, 95%.

**Supplementary Figure 4.**
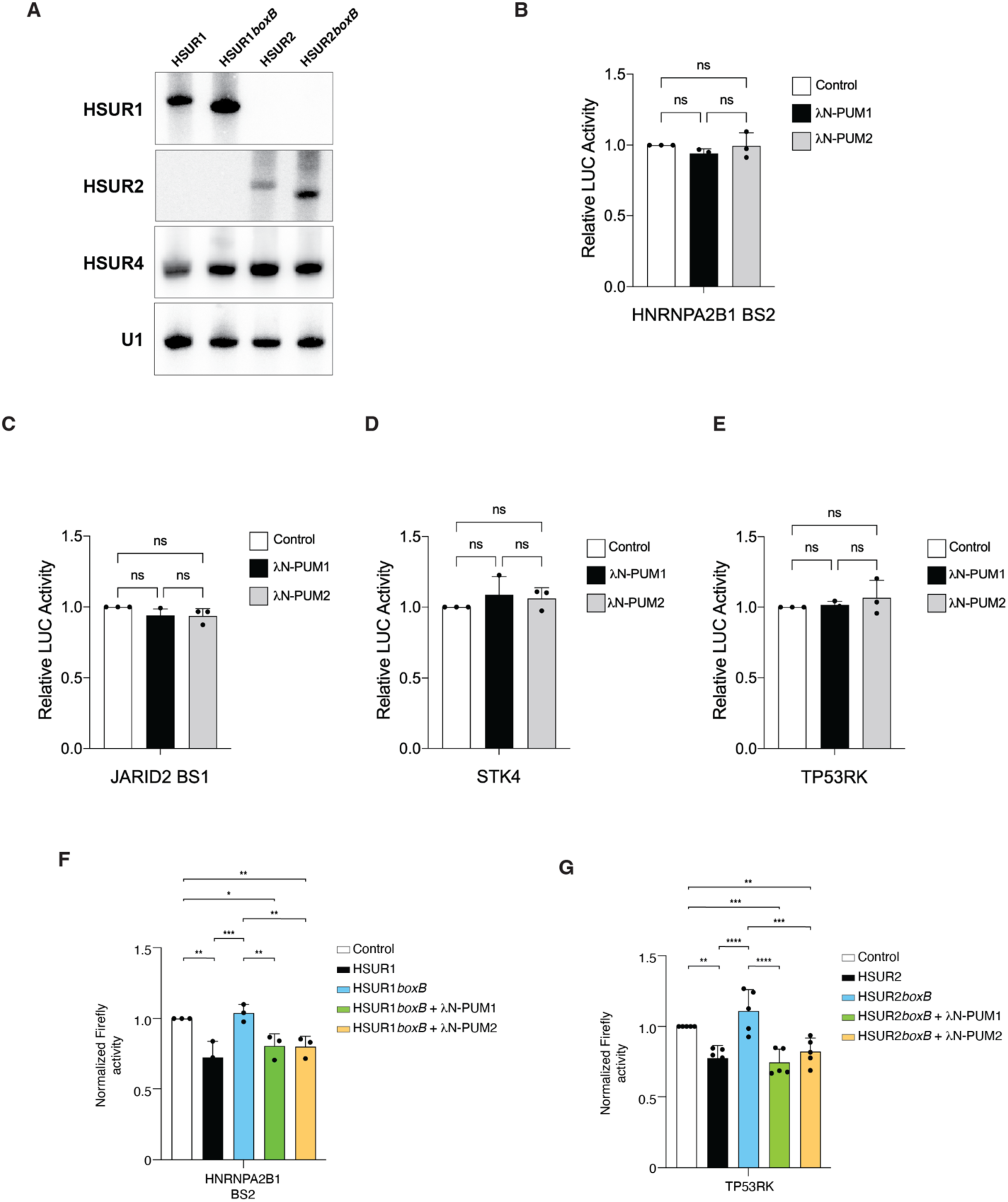
Association of PUM proteins with HSURs is required for HSUR-mediated mRNA repression. **(A)** Northen blot analyses of HeLa cells following transient co-transfection with a plasmid expressing HSUR4 and the plasmids used in Fig. 4 to express wild-type HSUR1 (143 nt), HSUR1*boxB* (137 nt), wild-type HSUR 2 (115 nt), or HSUR2*boxB* (107 nt). Membranes were probed for the indicated RNAs. HSUR4 served as a transfection control, whereas cellular U1 snRNA served as a loading control. **(B-E)** Luciferase assays in HuT78 cells stably expressing reporter transcripts containing 3′UTR fragments with single HSUR1-binding sites (B and C), or U937 cells stably expressing reporter transcripts containing 3′UTR fragments with single HSUR2-binding sites (D and E) following transient co-transfection of a control plasmid expressing mCherry2 alone (Control) or plasmids encoding mCherry2 and λN-PUM1 or λN-PUM2, together with a control plasmid containing the U1 promoter. Firefly luciferase levels were normalized to *Renilla* luciferase levels. **(F)** Same as in Fig. 4B with a reporter transcript containing a fragment of HNRNPA2B1 3′UTR with a single HSUR1 binding site (binding site 2, BS2). **(G)** Same as in Fig. 4C, using a reporter transcript containing a fragment of the TP53RK 3′UTR with a single HSUR2 binding site. \**P* < 0.05; \*\**P* < 0.01; \*\*\**P* < 0.001; \*\*\*\**P* < 0.0001.

**Supplementary Figure 5.**
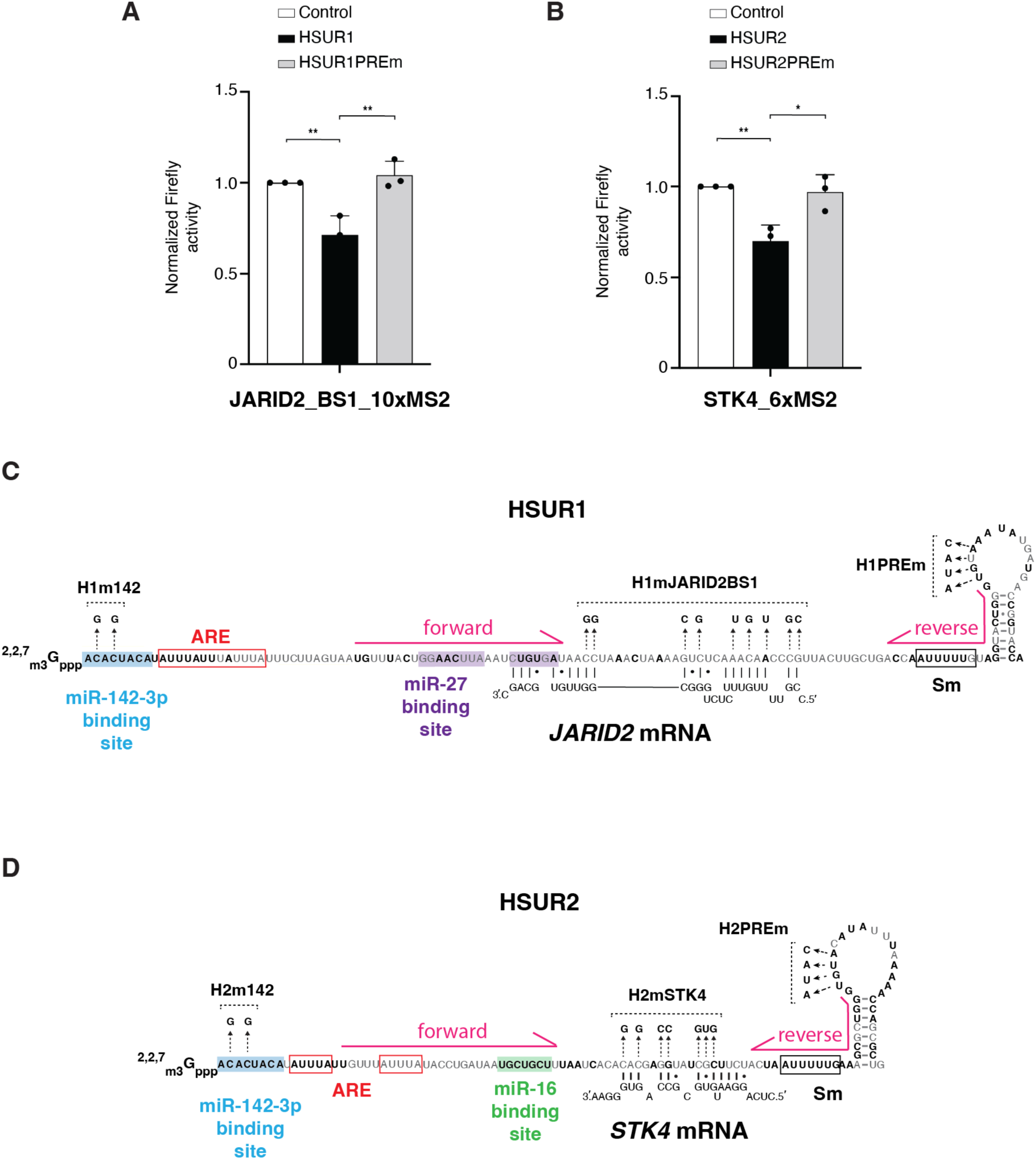
PUM proteins license HSUR association with target mRNAs. **(A)** Same as in Fig. 3C, using a reporter transcript containing a fragment of JARID2 3′UTR with a single HSUR1 binding site and ten MS2 hairpin repeats. **(B)** Same as Fig. 3D, using a reporter transcript containing a fragment of the STK4 3′UTR with a single HSUR2 binding site and six MS2 hairpin repeats. **(C)** Nucleotide sequence of HSUR1 showing the nucleotides modified in each mutant used in Fig. 5B. The position of the sequences complementary to the forward and reverse primers used to analyze HSUR1 enrichment performed in Fig. 5B are indicated by pink arrows. **(D)** Same as in (C) for HSUR2, showing the nucleotides modified in each of the mutants used in Fig. 5C and the primers used to analyze HSUR2 enrichment.

